# Saliva Metabolomics Reveals Distinct Metabolic Signatures in Patients with Chronic Obstructive Pulmonary Disease: A GC-MS-based approach

**DOI:** 10.64898/2026.04.10.717654

**Authors:** Rishita Singh, Sudip Ghosh, Neha Yadav, Amit Kumar Mandal

## Abstract

Chronic obstructive pulmonary disease (COPD), a chronic lung disease, involves complex metabolic disturbances that remain poorly characterized using non-invasive matrices. The metabolic alterations associated with cigarette smoke (CS), one of the major drivers of disease progression in COPD patients, have not been explored in detail. This study primarily aimed to investigate the metabolic signatures in COPD patients categorized into smoker (n=15), ex-smoker (n=11), and non-smoker (n=3) subgroups. Utilizing saliva as a noninvasive sample, we identified 26 metabolites with differential expression in smokers and 31 in ex-smokers. However, no such significant alteration was observed in the non-smokers subgroup. The multivariate analysis distinctly separated the COPD subgroups from healthy controls. Additionally, pathway enrichment analysis revealed perturbations in key metabolic pathways, including unsaturated fatty acid biosynthesis, arginine biosynthesis, the tricarboxylic acid (TCA) cycle, and pyruvate metabolism. Moreover, univariate Random forest analysis identified four metabolites (cyclopentanone, tetradecane 4-methyl, acetophenone, and scyllo-inositol) as potential biomarkers distinguishing COPD subgroups from healthy controls. This study offers novel molecular insights into the association of smoking with disease progression and provides a mechanistic understanding of COPD in different subgroups for better management of the disease.

**Graphical abstract:** 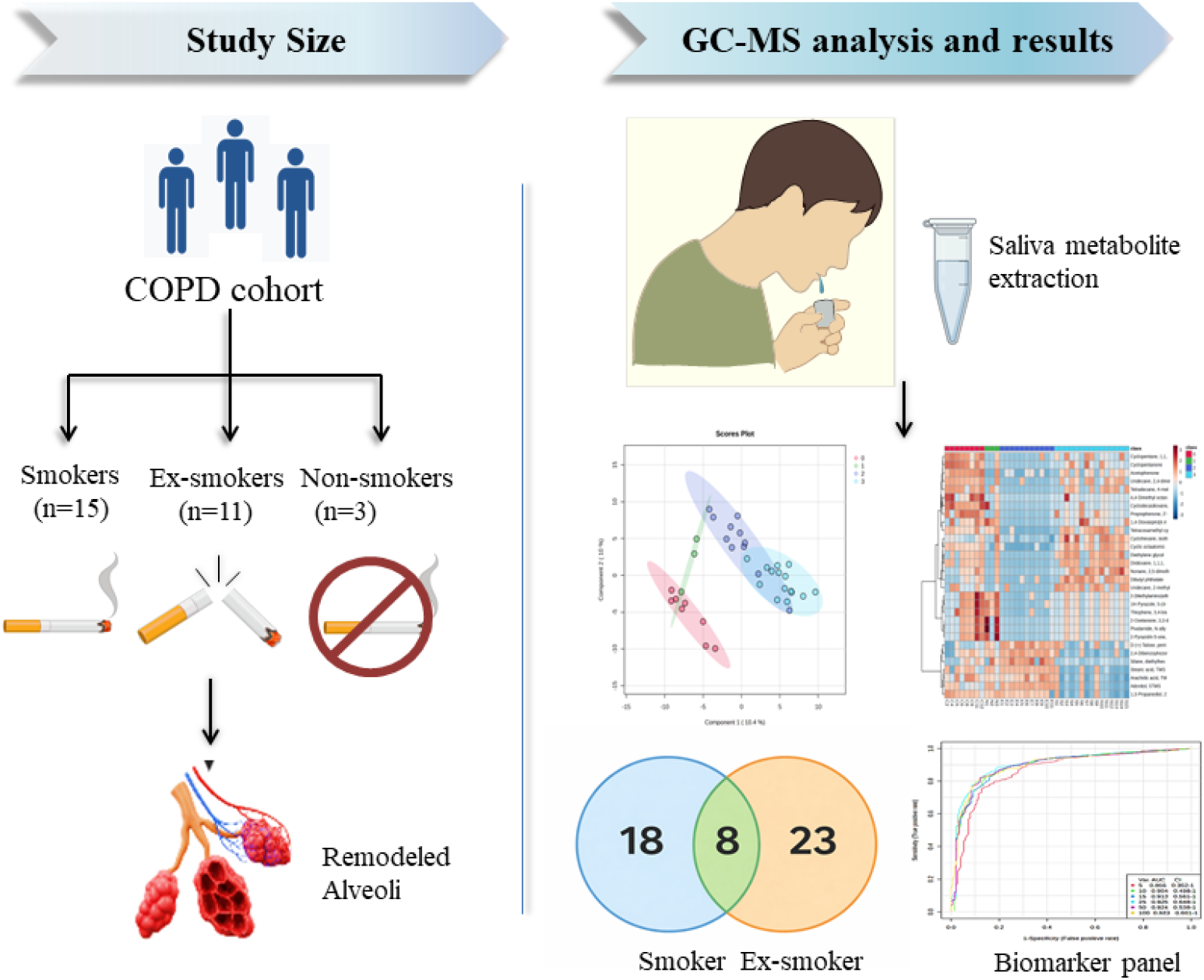

## 1. Introduction

Chronic Obstructive Pulmonary Disease (COPD) is a heterogeneous lung condition characterized by persistent respiratory symptoms (dyspnea, cough, sputum production) and exacerbations. These manifestations result from sustained, often progressive, airflow obstruction attributable to abnormalities in the airways (bronchitis, bronchiolitis) and/or alveoli (emphysema) **[1]**. It is the third major cause of morbidity and mortality across the globe, causing 3.7 million deaths worldwide annually. According to the Global Burden of Disease projection, COPD will remain the third leading cause of mortality by 2050 **[2]**. The number of COPD cases has increased to 213 million worldwide, with Asian countries having the highest number of COPD cases **[3]**. India has the highest burden of COPD cases, around 18% of total COPD cases, and around 27% of all COPD deaths globally **[4]**. In COPD, loss of alveolar wall elastic recoil leads to a decline in forced expiratory volume (FEV) and impaired lung emptying on expiration. It encompasses chronic bronchitis, bronchiolitis, emphysema, and pulmonary vascular remodelling, characterized by airway narrowing, parenchymal destruction, and structural changes in pulmonary vasculature. COPD is diagnosed using spirometry, with a post-bronchodilator FEV_1_/FVC<70% used as the diagnostic threshold **[5]**. Despite the availability of diagnostic tests, several barriers to accessing spirometry remain, including a lack of facilities in primary health care, limited awareness, and the risk of misdiagnosis, as reported in a study in which about 10% of doctor-diagnosed COPD were concordant with lung function results **[6]**. Furthermore, it is not uncommon for symptoms to appear very late in the disease stage **[7]**, and spirometry is not particularly sensitive in detecting early small airway obstruction. Additionally, a physiological decrease in FEV_1_/FVC occurs with age; therefore, the use of a fixed ratio (FEV_1_/FVC<70%) may contribute to overdiagnosis **[8]**.

There are several risk factors associated with COPD, like cigarette smoking, exposure to air pollutants originating from the burning of biomass and fossil fuels, and a deficiency of α1-antitrypsin. Among these, CS is one of the major risk factors for COPD, contributing 90% of total COPD cases **[9]**. CS comprises a complex mixture of reactive chemicals and oxidants that trigger oxidative stress, inflammation, metabolic dysregulation, etc, thereby accelerating the onset of disease and its progression. Although approximately 21% of cigarette smokers develop COPD, the disease is also observed in ex-smokers (∼10% of cases) and non-smokers (∼3.3% of cases) **[10]**. Despite the well-established role of smoking in COPD, the molecular mechanisms distinguishing COPD smokers, ex-smokers, and non-smokers remain poorly understood.

Recently, omics-based approaches, such as genomics, transcriptomics, proteomics, lipidomics, and metabolomics, have contributed to the understanding of several diseases. Metabolomic profiling offers unique insights into disease-associated biochemical changes, has strong potential for biomarker discovery, and may help elucidate COPD pathophysiology with respect to smoking and its role in exacerbation rates and disease severity. Previously, metabolomics studies in COPD have primarily focused on human biofluids, including sputum, plasma, and serum, and have reported perturbations in amino acid metabolism, tricarboxylic acid cycle intermediates, lipid remodeling, and redox imbalance **[11,12,13]**. However, there are no reports on the metabolic profile of saliva in COPD patients categorized into different subgroups based on the smoking history.

In this study, we performed the untargeted metabolomics of saliva samples using a GC-MS-based approach in patients with COPD stratified by smoking status, encompassing smokers, ex-smokers, and non-smokers subgroups. We evaluated and compared the distinct metabolic signatures associated with smoking, uncovering biochemical pathways that persist beyond smoking cessation and differentiate COPD subtypes at the molecular level.

## 2. Materials and methods

### 2.1. Materials

All chemicals of analytical grade, such as Methoxyamine hydrochloride (MOX), N-Methyl-N-(trimethylsilyl)trifluoroacetamide (MSTFA), Trimethylchlorosilane (TMCS), and Ribitol, were purchased from Sigma Aldrich (USA). LC-MS grade methanol, water, and HPLC-grade hexane and pyridine were purchased from Thermo Fisher Scientific (USA).

### 2.2. Study design

The study was conducted at the Indian Institute of Science Education and Research (IISER), Kolkata, in collaboration with All India Institute of Medical Sciences (AIIMS), Kalyani. The ethics committees of both institutions approved the study with reference no: IEC/AIIMS/Kalyani/certificate/2024/415. Participants with known cases of COPD per the GOLD standard (Global Initiative for Chronic Obstructive Lung Disease) were recruited after obtaining written informed consent. COPD patients with and without the habit of smoking were considered after a detailed smoking history assessment. Patients with post FEV_1_/FVC < 70% were included in the study, whereas participants with other comorbidities such as tuberculosis, asthma, diabetes, cancer, HIV (Human immunodeficiency virus), HBsAg (Hepatitis B surface antigen), and anti-HCV (Hepatitis C virus antibody) positive were excluded. Unstimulated saliva was collected post-spirometry from 29 COPD patients and 8 healthy controls. These COPD patients were further categorised into three different subgroups, i.e., smokers (n=15), ex-smokers (n=11), and non-smokers (n=3).

### 2.3. Sample collection

Approximately 2 mL of unstimulated saliva was collected in a plastic container using the drooling method, then centrifuged at 12,000 rpm for 20 minutes at 4 °C to remove insoluble mucus and debris. The supernatant obtained was then aliquoted and stored at -80 °C until further analysis.

### 2.4. Sample preparation

The stored saliva samples were thawed at 4 °C, and 200 μL of each sample was used for metabolite extraction. 10 μL of ribitol solution (0.4mg/mL) was added to each sample. 600 μL of a 3:1 (v/v) methanol: water mixture was used as the extraction solvent. After thorough mixing of the sample with the extraction solvent for 25 minutes at room temperature, the samples were centrifuged at 12,000 rpm for 30 minutes at 4 °C. The collected supernatant was vacuum-dried and then lyophilised. The lyophilised samples were reconstituted in 50 μL MOX solution (20 mg/mL) prepared in pyridine, vortexed for 30 seconds, and incubated overnight at 37 °C. A 50 μL of MSTFA containing 1% TMCS was then added to the samples, vortexed for 30 seconds, and again incubated at 80 °C for 1 hour and 30 minutes. A 50 μL of n-hexane was added to the sample, vortexed for 30 seconds, and finally centrifuged at 12,000 rpm for 10 minutes at 4 °C. The supernatant was collected for GC-MS analysis.

### 2.5. GC-MS Analysis

The prepared sample was analysed using an Agilent 8890 GC system coupled with a 7010C GC/Triple Quad MS system (Agilent Technologies, USA). The metabolite separation was performed on a DB-5MS column coated with 5% phenyl-methylpolysiloxane (30 m x 250 μm i.d., x 0.25 μm film thickness (Agilent Technologies, USA). The initial GC oven temperature was set to 60 °C for 1 minute, then increased at 10 °C/minute until the temperature reached 325 °C. The inlet, transfer line, and ion source temperatures were set at 250 °C, 290 °C, and 250 °C, respectively. The sample injection volume was 1 μL with a 1:5 split ratio; helium was used as the carrier gas at a flow rate of 1 mL/minute. The column temperature was set to 60 °C for 1 minute, then increased at 10 °C/minute to 325 °C and held for 10 minutes. Mass was acquired in full scan mode in the range 50-800 m/z, and ions were generated with electron impact ionization mode using 70 eV.

Saliva metabolites isolated from COPD patients in the following subgroup: smokers (n=15), ex-smokers (n=11), and non-smokers (n=3) were analysed in GC, and quality control (QC) for each category was run in between the sample batches to correct the variability during the run. For the comparative analysis with COPD patients, saliva metabolites isolated from healthy non-smoker subjects (n=8) were analysed.

### 2.6. Data Processing and Statistical Analysis

The raw data were analysed using Agilent MassHunter workstation software (Agilent Technologies, USA). Data pretreatment included baseline correction, deconvolution, retention-time alignment, peak filtering, and identification. The compound identities were annotated after spectral search with a match score ≥50 against the National Institute of Standards and Technology (NIST) library. Features with >50% missing values per group were removed, and the remaining missing values were imputed using one-fifth of the minimum positive value. The Pre-treated data were normalized using sum normalization, log-transformed, and Pareto-scaled using MetabolAnalyst 6.0 software. Multivariate analysis, including unsupervised Principal Component Analysis (PCA) and supervised Orthogonal Partial Least Squares Discriminant Analysis (OPLSDA), was performed to assess similarities and differences between groups. The OPLSDA model parameters R^2^Y (goodness of fit) and Q^2^ (predictive ability) were used to quantify the total data variance and the predictive power, respectively, with the p-value obtained from a permutation (n=2000). A univariate Wilcoxon rank-sum test with Benjamini-Hochberg correction was performed to assess the significant differences in metabolites between the subgroups, with FDR-adjusted p-values. The variable importance in projection (VIP score >1), fold change (FC ≥2), p-value <0.05, and false discovery rate (FDR <0.05) were all taken into consideration to obtain differential metabolites between each subgroup of COPD and the control group.

### 2.7. Biomarker analysis

Random forest analysis was conducted to identify the potential biomarker. Metabolites with FDR-adjusted p value <0.05, VIP >1, and FC ≥2 were taken for biomarker analysis. The diagnostic potential of these metabolites was assessed using receiver operating curve (ROC) analysis. Metabolites with an area under the curve (AUC) >0.8 were considered to possess good diagnostic ability.

### 2.8. Pathway enrichment and correlation analysis

Pathway enrichment analysis was performed using Kyoto Encyclopedia of Genes and Genomes (KEGG) and the Human Metabolome Database (HMDB) in MetaboAnalyst 6.0 to identify metabolic pathways associated with altered metabolites in COPD subgroups. The differentially expressed metabolites obtained after VIP score >1, p-value <0.05, and FC ≥2 were subjected to pathway enrichment analysis. Spearman’s rank correlation analysis was performed to assess the association between differentially expressed metabolites in COPD subgroups and spirometry index, i.e., post FEV_1_/FVC ratio, with a correlation coefficient (ρ ≥ 0.3) and p-value < 0.05.

## 3. Results

### 3.1. General Characteristics of the participants

The study includes 29 COPD patients classified into the following subgroups: 15 COPD smokers, 11 COPD ex-smokers, 3 COPD non-smokers, and 8 healthy controls. Spirometry data and vital signs from patients and healthy individuals were analysed and reported as mean values with standard deviations. The statistical parameters are obtained using GraphPad Prism 8.0.2. Detailed characteristics of the patients and control groups are given in Table 1.

**Table 1.** Clinical parameters of COPD patients and healthy controls.

| Patient's clinical details | COPD patient<br>Smoker<br>(n=15) | COPD patient<br>Ex-smoker<br>(n=11) | COPD patient<br>Non-smoker<br>(n=3) | Healthy<br>Non-smoker<br>(n=8) |
| --- | --- | --- | --- | --- |
| Age | 60.60 ± 8.41 | 61.85 ± 8.99 | 49.67 ± 17.2 | 25.63 ± 3.15 |
| Sex (M/F) | 15/0 | 11/0 | 1/2 | 6/2 |
| Height in cm | 159.4 ± 6.15 | 162.3 ± 4.94 | 153.7 ± 2.0 | 166.3 ± 6.45 |
| Weight in Kg | 54.0 ± 10.73 | 61.45 ± 10.87 | 48.0 ± 1.7 | 67.18 ± 14.9 |
| BMI | 21.22 ± 3.65 | 23.45 ± 4.56 | 20.3 ± 1.2 | 24.7 ± 4.7 |
| Cigarette (Pack years) | 42.95 ± 23.87 | 25.3 ± 27.46 | N/A | N/A |
| Smoking years | 34.07 ± 8.46 | 24.17 ± 13.05 | N/A | N/A |
| Pre-FEV1 | 0.93 ± 0.40 | 0.81 ± 0.32 | 1.13 ± 0.30 | 3.06 ± 0.73 |
| Post- FEV1 | 1.11 ± 0.41 | 0.86 ± 0.32 | 1.20 ± 0.32 | 3.18 ± 0.75 |
| Pre- FVC | 2.06 ± 0.63 | 1.9 ± 0.67 | 2.42 ± 0.66 | 3.59 ± 0.96 |
| Post- FVC | 2.26 ± 0.69 | 2.04 ± 0.74 | 2.34 ± 0.69 | 3.6 ± 0.95 |
| Pre- FEV1/FVC | 44.88 ± 10.8 | 45.3 ± 18.3 | 46.80 ± 1.35 | 85.58 ± 5.4 |
| Post- FEV1/FVC | 48.6 ± 15.46 | 44.7 ± 16.9 | 51.53 ± 4.31 | 88.3 ± 5.0 |
| All data points are Mean ± SD, n = number of samples, FEV1 = Forced expiratory volume in 1 sec, FVC = Forced vital capacity |  |  |  |  |

**Table 2.** List of four significantly altered metabolites across COPD subgroups (smoker and ex-smoker) in comparison to healthy controls.

| S.No | Metabolites | HMDB | p-value | VIP | FDR | AUC |
| --- | --- | --- | --- | --- | --- | --- |
| 1 | Cyclopentanone | HMDB0031407 | 9.92E-06 | 1.5889 | 0.001935 | 0.952586 |
| 2 | Acetophenone | HMDB0033910 | 0.000161 | 1.6115 | 0.002843 | 0.965517 |
| 3 | Tetradecane<br>4-methyl | HMDB0302457 | 2.82E-05 | 1.2777 | 0.001959 | 0.909483 |
| 4 | Scyllo-Inositol | HMDB0006088 | 0.007745 | 1.0824 | 0.033192 | 0.896552 |

### 3.2. Comparative metabolic Profiling between different COPD subgroups

Upon analyzing salivary GC-MS metabolomics data using MetaboAnalyst 6.0, both unsupervised PCA and supervised Partial Least Squares Discriminant Analysis (PLS-DA) separated the COPD subgroups (smokers, ex-smokers, and non-smokers) from healthy controls **(Fig 1A, B)**. The cross-validated PLS-DA model yielded acceptable performance with R^2^ = 0.79 and Q^2^ = 0.49, indicating differentiation among the COPD subgroups. Univariate analysis using a nonparametric Kruskal-Wallis test, with Benjamini-Hochberg, identified 107 significantly altered metabolites with FDR-corrected p-value <0.05. These metabolites were further refined using VIP score >1 and AUC > 0.8, resulting in the identification of 13 metabolites **(Table S1, supplementary file)**. The VIP score obtained from PLS-DA is depicted in **Fig. 1C**, and the hierarchical clustering heatmap is given in **Fig S6**. Furthermore, to assess pairwise comparisons, a Wilcoxon rank-sum test with the Benjamini-Hochberg correction was performed.

**Fig. 1.**
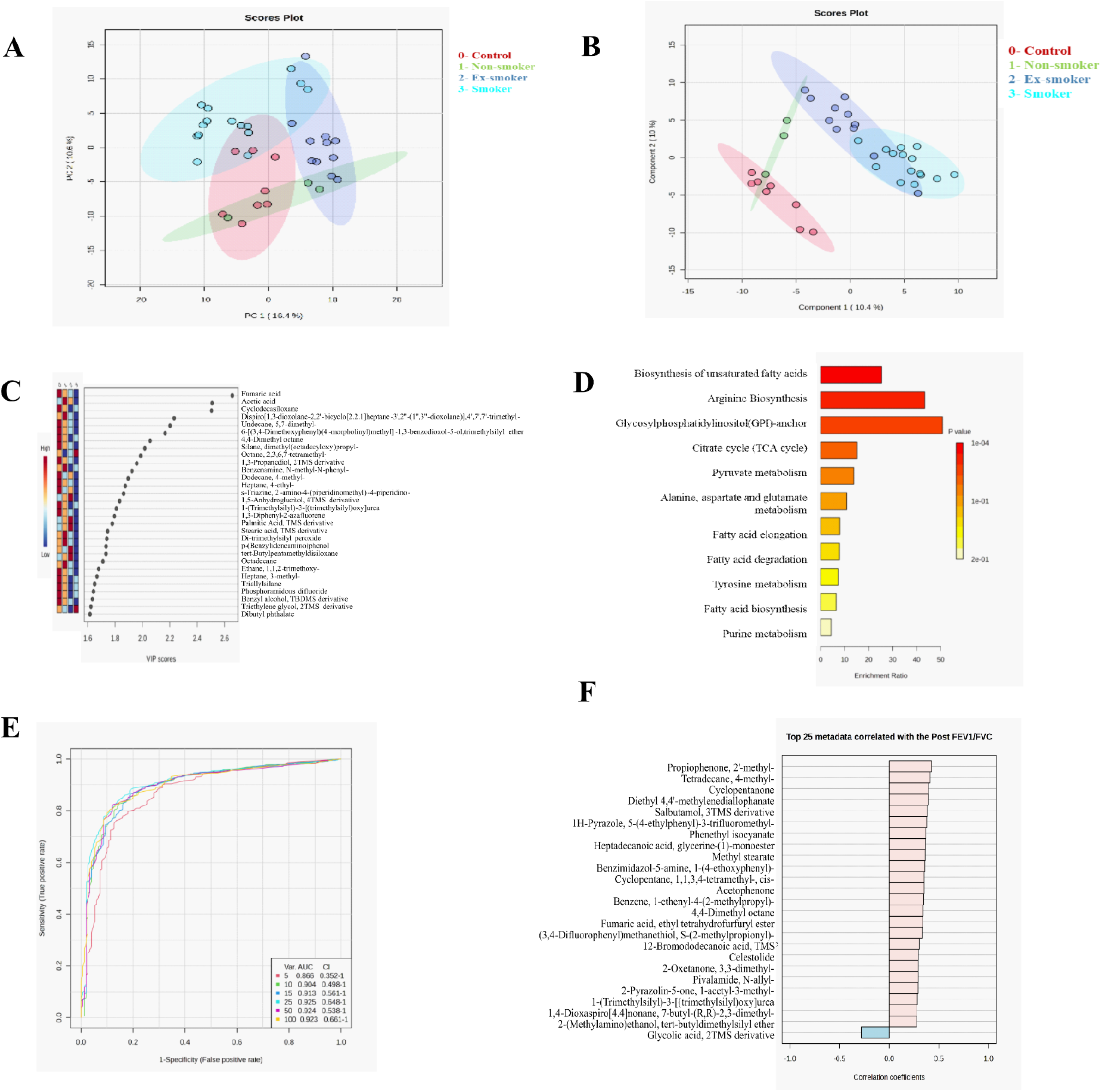
The comparison of saliva metabolites between various COPD subgroups and healthy controls is illustrated as follows: (A) A PCA score plot differentiates COPD subgroups from healthy controls, with green representing COPD non-smokers, blue indicating COPD ex-smokers, cyan denoting COPD smokers, and red signifying healthy controls. (B) A PLSDA score plot further separates the respective subgroups, (C) A discriminant analysis VIP score plot highlights differential metabolites, (D) An overview of pathway enrichment reveals dysregulated pathways in COPD subgroups, (E) A multivariate ROC curve, generated using a random forest algorithm, represents all models. (F) Spearman’s correlation is depicted between significant metabolites and post FEV_1_/FVC.

Multivariate analysis comparing COPD smokers with healthy controls revealed distinct group separation through PCA and OPLSDA analysis **(Fig 2A, B)**. The OPLSDA model exhibited a robust goodness-of-fit (R^2^Y = 0.88) and predictive ability (Q^2^ = 0.80) with statistical validation using permutation testing (n = 2000), yielding R^2^Y = 0.98 (p-value = 0.0015), Q^2^ = 0.88 (p-value = 5E-04), respectively. A total of 26 significant metabolites were identified, with VIP scores >1, p-value <0.05, FC ≥2, FDR <0.05, and AUC >0.8 **(Table S2 and Fig S1)**. The corresponding volcano plot with p-values and log_2_FC, alongside the VIP score plot, is depicted in **Fig 2C and 2D**, respectively. The hierarchical clustering heatmap is represented in **Fig S7**.

**Fig 2.**
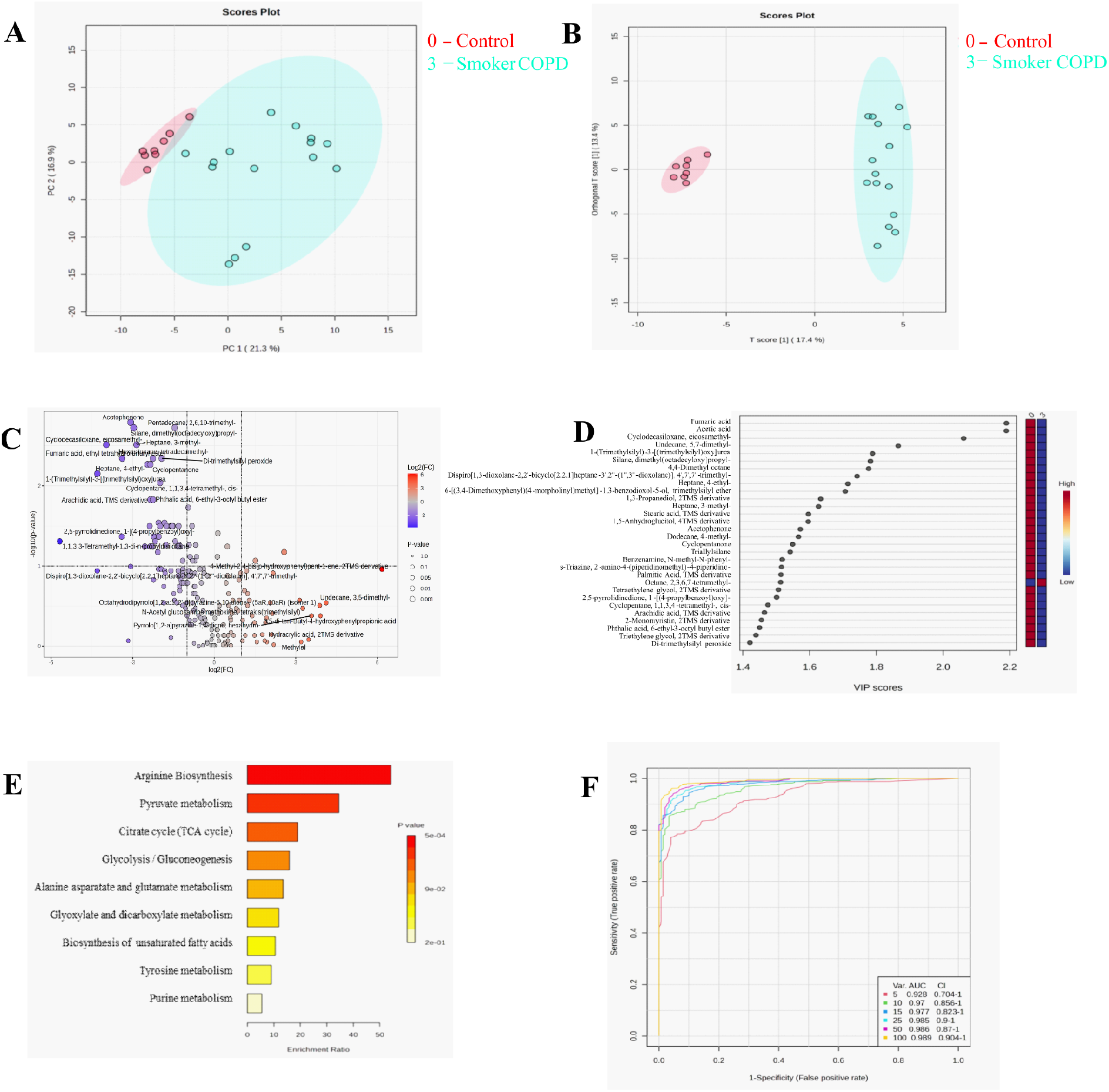
(A) PCA score plot differentiating COPD smokers (cyan) from healthy controls (red), (B) OPLSDA score plot of COPD smokers (cyan) and healthy controls (red), (C)Volcano plot showing differentially expressed metabolites between smokers vs healthy controls, (D) Discriminant analysis VIP score plot of differential metabolites, (E) Pathway enrichment overview of dysregulated pathways in smokers, (F) Multivariate ROC curve generated from random forest algorithm representing all models.

PCA and OPLSDA analysis revealed a distinct separation between COPD ex-smokers and healthy controls **(Figure 3A, B)**, with R^2^Y = 0.94 and Q^2^ = 0.86. The robustness of the model was confirmed through permutation testing (n=2000), yielding R^2^Y=0.99 (p-value =5E-04) and Q^2^ =0.89 (p-value =5E-04), indicating significant statistical differentiation between the groups. Wilcoxon rank sum test identified 55 metabolites with FDR-adjusted p-values <0.05, which were subsequently filtered using VIP score >1, FC ≥2, and AUC >0.8, resulting in a total of 31 metabolites **(Table S3, and Fig S2)**. The volcano plot and the VIP score plot are presented in **Fig 3C and 3D**, respectively. The hierarchical clustering heatmap is shown in **Fig S8**.

**Fig 3.**
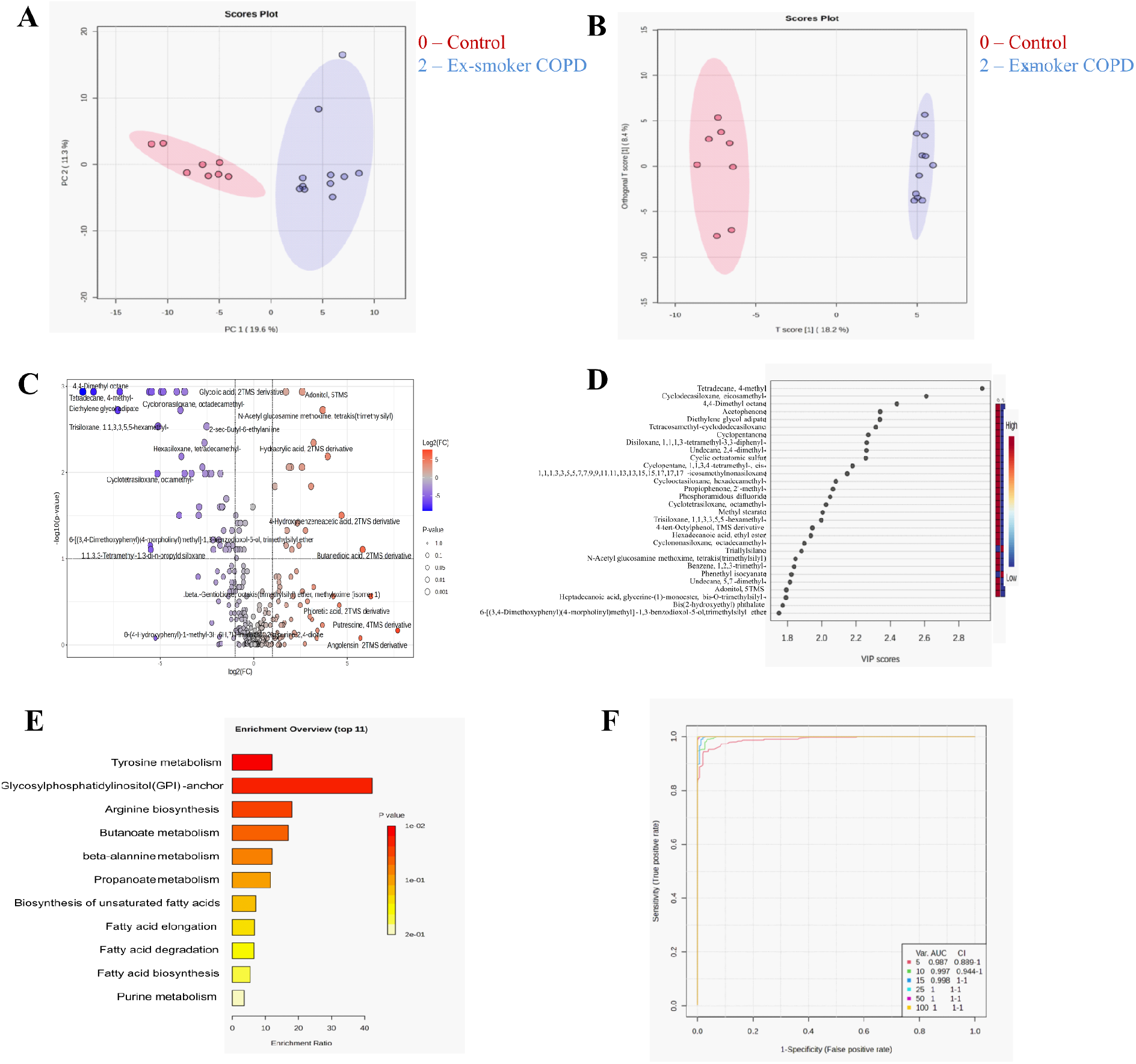
(A) PCA score plot comparing COPD ex-smokers (blue) with healthy controls (red), (B) OPLSDA score plot of COPD ex-smokers (blue) and healthy controls (red), (C)Volcano plot illustrating differentially expressed metabolites between ex-smokers vs healthy control, (D) Discriminant analysis VIP score plot of differential metabolites, (E) Pathway enrichment overview highlighting dysregulated pathways in ex-smokers, (F) Multivariate ROC curve generated from random forest algorithm representing all models.

Additionally, PCA and OPLSDA analysis of COPD non-smokers versus healthy controls showed no clear separation between groups **(Fig 4A, B)**, with R^2^Y = 0.7 and Q^2^ = 0.24. Furthermore, permutation validation (n=2000) confirmed a nonsignificant difference with R^2^Y=0.99 (p-value=0.22), and Q^2^ =0.40 (p-value=0.16), respectively. Upon performing a Wilcoxon rank-sum test, no significant difference was found between the groups, further supporting the multivariate findings. The compounds, such as decasiloxane, triallyl silane, etc., have been identified as GC column contaminants and were not processed for downstream analysis.

**Fig 4.**
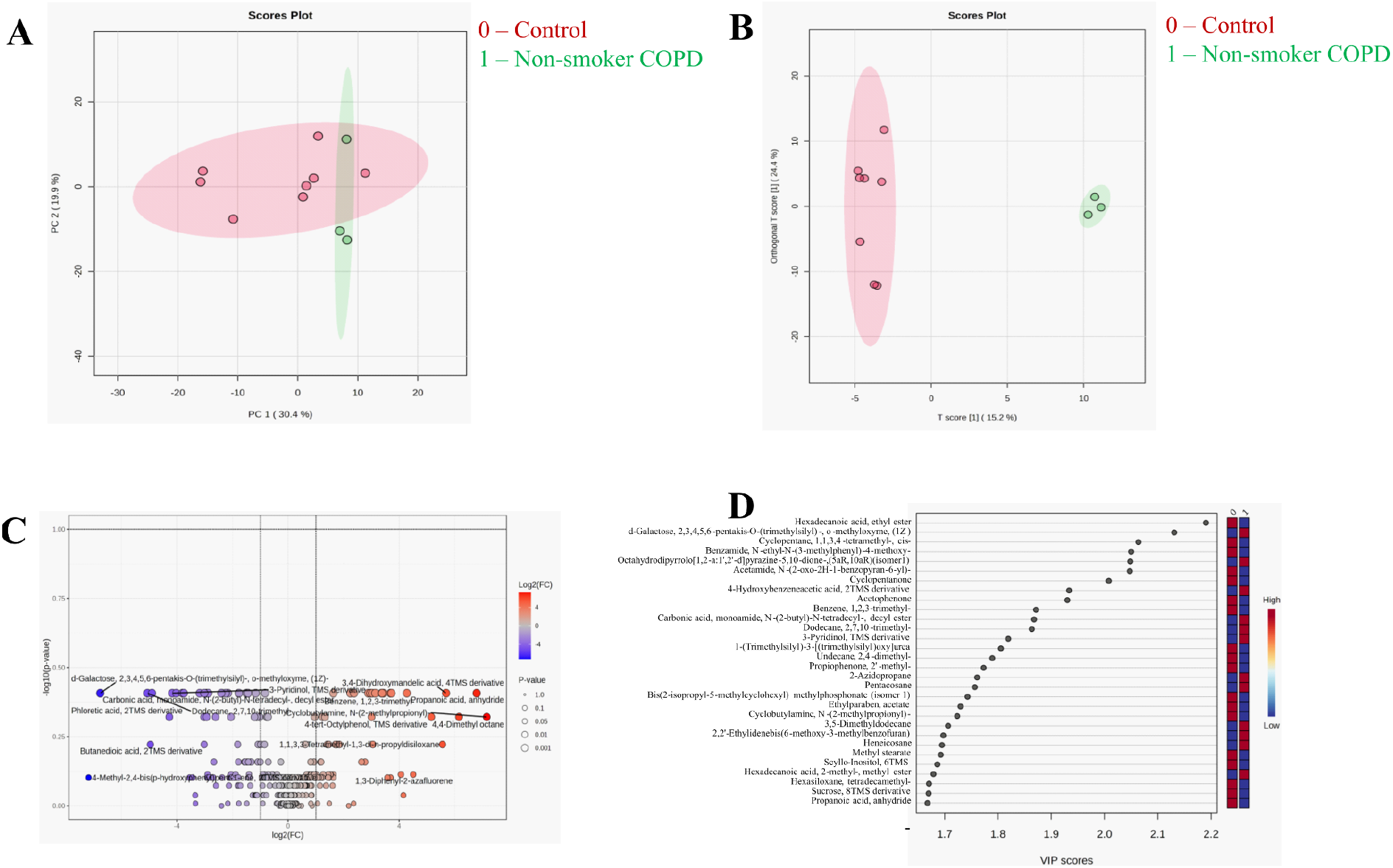
(A) PCA score plot comparing COPD non-smokers (green) with healthy controls (red), (B) OPLSDA score plot contrasting COPD non-smokers (green) and healthy controls (red), (C) Volcano plot indicating no significant differential expression of metabolites between non-smokers and healthy controls, (D) Non-robust discriminant analysis VIP score plot.

**Fig 5.**
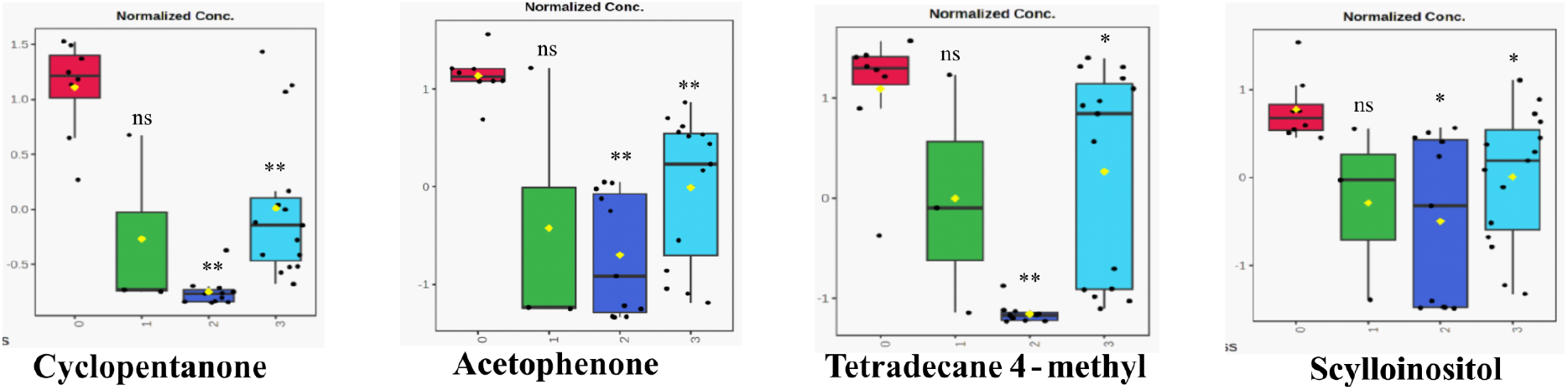
Box and whisker plots of four differentially expressed metabolites across all COPD subgroups (cyan: smokers, blue: ex-smokers, green: non-smokers, red: healthy controls) with *: p< 0.05, **: p< 0.01, ns: non significant.

**Fig 6:**
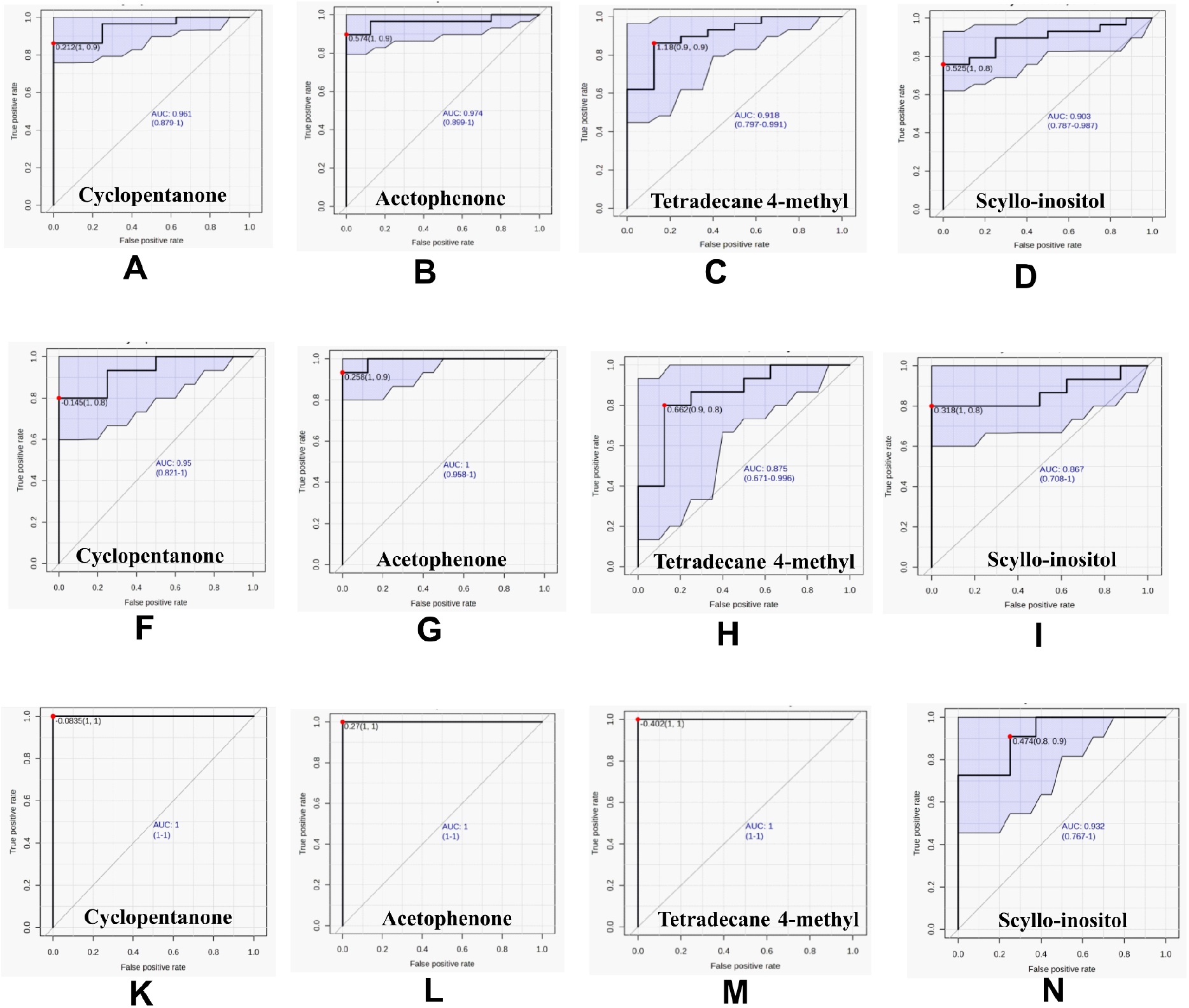
The receiver operating characteristic (ROC) curves of four differential metabolites, each with an area under the curve (AUC) of ≥ 0.85, are presented for various comparison groups: (A-D) COPD subgroups versus healthy controls, (F-I) COPD smokers versus healthy controls, and (K-N) COPD ex-smokers versus healthy controls.

### 3.3. Pathway analysis

Pathway enrichment analysis of differentially abundant metabolites across all COPD subgroups versus healthy controls identified significant perturbations in metabolic pathways, including unsaturated fatty acid biosynthesis, arginine biosynthesis, glycosylphosphatidylinositol (GPI)-anchor biosynthesis, and TCA cycle **(Fig 1D)**. Comparison of significant metabolites between the COPD smokers subgroup and healthy controls highlighted disruptions in arginine biosynthesis, pyruvate metabolism, the TCA cycle, and glycolysis/gluconeogenesis **(Fig 2E)**. Comparison of significantly altered metabolites between COPD ex-smokers and healthy controls revealed perturbations in pathways including tyrosine metabolism, GPI-anchor biosynthesis, arginine biosynthesis, and butanoate metabolism **(Fig 3E)**. In summary, the major perturbed pathways encompassed fatty acid, amino acid, and energy metabolism.

### 3.4. Biomarker analysis

The ROC curve obtained from Random Forest analysis identified significant features that discriminate COPD subgroups from healthy controls. Features with AUC>0.8, (95% CI (confidence interval)), p-value <0.05, and FC ≥2 are considered to be a potential diagnostic marker for COPD subgroups. Four metabolites have been consistently identified across all comparisons (COPD vs. controls, COPD smokers vs. controls, and COPD ex-smokers vs. controls). These include cyclopentanone with AUC value 0.96 (95% CI: 0.87-1); 0.95 (95% CI: 0.82-1); 1 (95% CI: 1-1), acetophenone with AUC 0.97 (95% CI: 0.89-1); 1 (95% CI: 0.95-1); 1 (95% CI: 1-1), tetradecane 4-methyl with AUC 0.91 (95% CI: 0.79-0.99); 0.87 (95% CI: 0.67-0.99); 1 (95% CI: 1-1), and scyllo-inositol with AUC 0.90 (95% CI: 0.78-0.98); 0.86 (95% CI: 0.7-1); 0.93(95% CI: 0.76-1), for COPD vs healthy control; COPD smoker vs healthy control; and COPD ex-smoker vs healthy control, respectively (**Fig 7)**. Boxplots for these metabolites are shown in **Fig 8**.

However, groupwise comparison reveals subgroup-specific features with strong AUC and 95% CI values. In COPD smokers vs healthy control: pentadecane 2,6,10-trimethyl-(AUC= 0.98, 95%CI: 0.9-1), heptane 3-methyl (AUC= 0.96, 95% CI: 0.83-1), fumaric acid (AUC= 0.95, 95% CI: 0.82-1) were identified **(Fig S3)**. While in COPD ex-smokers vs healthy control, glycolic acid (AUC= 1, 95% CI: 1-1), N-acetyl glucosamine (AUC= 0.98, 95% CI: 0.9-1), and undecane 2,4-dimethyl (AUC= 1, 95% CI: 0.95-1) were detected **(Fig S3)**.

### 3.5. Correlation analysis

Spearman’s rank correlation analysis identified metabolites associated with spirometric post-bronchodilator FEV_1_/FVC ratio. COPD subgroups shows positive correlation with cyclopentanone (ρ = 0.39), acetophenone (ρ = 0.35), tetradecane 4-methyl (ρ = 0.41), propiophenone 2-methyl (ρ = 0.42), methyl stearate (ρ = 0.36), fumaric acid (ρ = 0.34), heptadecanoic acid (ρ = 0.36), and phenethyl isocyanate (ρ = 0.36) with p-value < 0.05 and negative correlation with glycolic acid (ρ = **-**0.28) and 3-hydroxybutyric acid (ρ = **-**0.23) with p-value > 0.05 **(Fig 1F)**. COPD smokers revealed positive correlation with heptane 4-ethyl (ρ = 0.56), fumaric acid (ρ = 0.48), and urea (ρ = 0.44) with p-value <0.05 and negative correlation with methyl-2,4-dihydroxy-3-methylbenzoate (ρ = **-**0.3), Bis(2-isopropyl-5-methylcyclohexyl) methylphosphonate (isomer1) (ρ = **-**0.22) with p-value >0.05 **(Fig S4)**. In comparison, COPD ex-smokers subgroup presented positive correlation with cyclopentanone (ρ = 0.7), acetophenone (ρ = 0.72), undecane 2,4-dimethyl (ρ = 0.69), tetradecane 4-methyl (ρ = 0.68), hexadecanoic acid ethyl ester (ethyl palmitate) (ρ = 0.61), propiophenone 2-methyl (ρ = 0.58), and methyl stearate (ρ = 0.51) with p-value <0.05 and negative correlation with hexadecane (ρ = **-**0.24), thiophene (ρ = **-**0.23) and acenaphthene (ρ = **-**0.12) with non significant p-value >0.05 **(Fig S5)**.

## 4. Discussion

COPD, primarily caused by cigarette smoking, exhibits a distinct metabolic profile **[11,12,14]**. However, metabolic differences across COPD subgroups, such as smokers, ex-smokers, and non-smokers, have not been profiled in saliva. Therefore, we investigated the subgroup-specific salivary metabolome and identified several altered classes of metabolites in COPD subgroups. In this study, we compared individual subgroups with healthy controls and identified altered metabolites within each subgroup. We observed significant alteration in the metabolomic profile of smokers and ex-smokers COPD patients compared to the healthy controls, whereas no significant alterations were observed in the COPD non-smoker subgroup. This highlights the role of smoking in disease stratification in COPD patients.

Compared to the healthy control, the smokers subgroup with COPD exhibited a significant decrease in the levels of metabolites such as fumaric acid, acetic acid, urea, etc. Fumaric acid, an unsaturated fatty acid involved in the TCA and urea cycles, indicates disruptions in energy metabolism and nitrogen handling, consistent with Hara *et al*.*’*s findings that CS induced oxidative stress in COPD, impairs mitochondrial function, and the TCA cycle **[15]**. The decreased urea level suggests disruption of the urea cycle, as previously reported in chronic liver disease and oral squamous cell carcinoma (OSCC) **[16,17]**. Similarly, reductions in fumaric acid and urea have been reported in cerebrovascular diseases, including Alzheimer’s disease and moyamoya disease **[18,19]**. The decline in acetic acid, a gut microbiota-derived short-chain fatty acid central to acetyl-CoA production and energy metabolism, aligns with studies linking its decrease to increased risk of bronchopulmonary dysplasia **[20]**. In addition, we observed a significant reduction in scyllo-inositol and arachidic acid levels in the subgroup of smokers with COPD. Scyllo-inositol is known to play a crucial role in protection against neurodegeneration **[21]**. The decrease in arachidic acid, a long-chain saturated fatty acid derived from arachidonic acid and associated with neuroprotection in autism spectrum disorder (ASD) **[22,23]**, is consistent with its downregulation in the plasma of COPD patients **[12]**. This might suggest that there is a possibility of neurodegeneration in the smokers subgroup of COPD patients **[24]**. The pathway analysis consolidates these findings by revealing broad perturbations in energy metabolism, fatty acid, and amino acid biosynthesis pathways, underscoring systemic metabolic dysregulation in smokers with COPD.

Compared with healthy controls, the ex-smoker COPD subgroup showed upregulation of glycolic acid, 3-hydroxybutyric acid, N-acetylglucosamine, and 4-hydroxybenzeneacetic acid. Glycolic acid and 3-hydroxybutyric acid have been identified as OSCC biomarkers **[17]**, whereas elevated N-acetylglucosamine is linked to type 2 diabetes **[25]**. The gut microbial metabolite 4-hydroxybenzeneacetic acid is associated with colorectal cancer **[26]**. Additionally, downregulation of scyllo-inositol, hexadecanoic acid ethyl ester (ethyl palmitate), and methyl stearate (stearic acid methyl ester) were observed. Hexadecanoic acid ethyl ester, known for its anticancer, anti-inflammatory, antimicrobial, and antioxidant effects, was decreased, possibly due to elevated oxidative stress. Hexadecanoic acid has been previously reported to be downregulated in COPD patients **[12,27]**. Methyl stearate, a saturated fatty acid, exhibits a neuroprotective and inhibitory role against the detrimental effects of cardiac arrest **[28]**. It has also been reported to play a role in the repair of cartilage defects **[29]**. Further, pathway analysis confirms the findings by revealing perturbations in fatty acid and amino acid metabolism.

We observed alterations in various volatile organic compounds (VOCs) in both smoker and ex-smoker subgroups, arising from the metabolic oxidation of macromolecules **[14]**; consequently, their concentrations may vary due to disease **[30]**. In this study, a reduction in the VOC levels was observed, possibly due to oxidative fragmentation of longer-chain hydrocarbons under elevated oxidative stress **[30]**. Undecane 4,7-dimethyl has previously been reported as a breath marker for distinguishing COPD from healthy controls **[30]**. In our findings, a notable decrease in undecane 5,7-dimethyl levels was observed in the subgroup of smokers with COPD. Additionally, we found reduced levels of undecane 2,4-dimethyl and undecane 2,6-dimethyl in the subgroup of ex-smokers. In this study, cyclopentanone, tetradecane 4-methyl, and acetophenone were observed to be downregulated. Previously, cyclopentanone and tetradecane were reported to be downregulated in breast cancer **[31]**, while tetradecane was recognized as a breath marker in discriminating obstructive sleep apnoea from obese patients **[32]**. Acetophenone has been reported as a potential biomarker in breast cancer and malignant pleural mesothelioma **[33,34,35,36]**.

These altered metabolites showed positive and negative correlation with post FEV_1_/FVC ratio. Additionally, random forest analysis identified 4 biomarker panels that are common to the smokers and ex-smokers subgroups. These four shared biomarkers, such as cyclopentanone, tetradecane 4-methyl, acetophenone, and scyllo-inositol, offer high specificity for smoking-related COPD, distinguishing smokers and ex-smokers from the non-smoker COPD subgroup and healthy controls. In the future, this biomarker panel could be validated in a larger COPD cohort to establish it as a noninvasive diagnostic marker, integrated with spirometry for early point-of-care use. These altered metabolites can again be used for targeted therapeutic interventions. These findings indicate a strong association between COPD and CS, suggesting disruption in energy and nitrogen metabolism, and probable elevation in the risks of cardiovascular dysfunction, neurodegeneration, and oral cancer development in the COPD smokers subgroup. At the same time, ex-smokers retain CS related metabolic signatures despite cessation. In contrast, the non-smokers subgroup of COPD showed no significant metabolic alteration as reflected in the smokers and ex-smokers subgroups.

## 5. Conclusion

The findings of our study highlight the clinical significance of saliva as a sample for COPD metabolomics, offering novel molecular insights into smoking-associated metabolic changes in COPD subgroups. In this study, through ROC and correlation analysis, we identify four potential biomarkers specific to the smokers and ex-smokers COPD subgroup: cyclopentanone, tetradecane 4-methyl, acetophenone, and scyllo-inositol. To further validate these biomarker panels, a large cohort study is essential. Overall, this work enhances our understanding of the metabolic pathways and mechanisms that contribute to the increased disease severity in COPD patients in the context of smoking. Thereby, advancing the development of potential non-invasive biomarkers and improved molecular phenotyping of COPD.

## 6. Ethics Declaration

This study was approved by the ethics committee of IISER Kolkata and AIIMS Kalyani (India) with reference number: IEC/AIIMS/Kalyani/certificate/2024/415

## 7. Author contribution

RS and AKM did the conceptualization. SG provided the samples. AKM and SG supervised the study and provided the resources and facilities. RS designed the study, collected the sample and demographic details of the participants, performed all the experiments, analysed data, wrote the original draft, reviewed and edited the manuscript. NY contributed to data analysis and was involved in writing the original draft and editing the manuscript. AKM reviewed and edited the manuscript.

## 8. Conflicts of interest

The authors declare no conflict of interest.

## 9. Acknowledgements

The author acknowledges the Council of Scientific and Industrial Research (CSIR), Government of India, for providing the fellowship. We would like to acknowledge all the participants who provided samples for this study, and Namrata Yadav (AIIMS-Kalyani) for helping in sample collection. We acknowledge the Nano Mission, Department of Science and Technology, for funding a mass spectrometry facility sanctioned under the project SR/NM/NS-1068/2015. We thank the DST-FIST (Project no.: SR/FST/LS-II/2017/93) for the common instrument facility at IISER Kolkata. We would also like to acknowledge the Department of Earth Science (DES) at IISER Kolkata for providing access to the GC-MS instrument.

## Notes

### Competing Interest Statement

The authors have declared no competing interest.

