## Supplemental data for "Saliva Metabolomics Reveals Distinct Metabolic Signatures in Patients with Chronic Obstructive Pulmonary Disease: A GC-MS-based approach"

### Table of contents:

#### Supplemental Tables

**Supplementary Table S1:** List of significantly altered metabolites between COPD subgroups and healthy controls.

**Supplementary Table S2:** List of significantly altered metabolites between COPD smokers and healthy controls.

**Supplementary Table S3:** List of significantly altered metabolites between COPD ex-smokers and healthy controls

#### Supplemental Figures

**Supplementary Figure S1:** Box and whisker plots of differentially expressed metabolites in COPD smokers vs healthy controls.

**Supplementary Figure S2:** Box and whisker plots of differentially expressed metabolites in COPD ex-smokers vs healthy controls.

**Supplementary Figure S3:** Receiver operating curves (ROC) of metabolites in smokers and ex-smokers subgroup.

**Supplementary Figure S4:** Correlation analysis of post $FEV_1/FVC$ ratio with differentially expressed metabolites in COPD smokers subgroups.

**Supplementary Figure S5:** Correlation analysis of post $FEV_1/FVC$ ratio with differentially expressed metabolites in COPD ex-smoker subgroups.

**Supplementary Figure S6:** Heatmap showing differential regulation of metabolites in COPD subgroups vs healthy controls.

**Supplementary Figure S7:** Heatmap showing differential regulation of metabolites in the COPD smoker subgroup vs healthy control.

**Supplementary Figure S8:** Heatmap depicting differential regulation of metabolites in the COPD ex-smoker subgroup vs healthy control.

**Table S1:** List of significantly altered metabolites between COPD subgroups and Healthy controls with VIP >1, p-value <0.05, FDR <0.05, and AUC>0.8.

| S.No. | Metabolites | HMDB ID | p-value | VIP | FDR | AUC |
|---|---|---|---|---|---|---|
| 1 | Cyclopentanone | HMDB0031407 | 0.00000992 | 1.5889 | 0.001935 | 0.962586 |
| 2 | Propiophenone 2-methyl | | 0.0000206 | 1.2822 | 0.001959 | 0.910862 |
| 3 | Tetradecane 4-methyl | HMDB0302457 | 0.0000282 | 1.2777 | 0.001959 | 0.914483 |
| 4 | 4,4 dimethyl octane | | 0.000037 | 2.0558 | 0.001959 | 0.887931 |
| 5 | Cyclopentane | | 0.0000603 | 1.4219 | 0.001959 | 0.943966 |
| 6 | Acetophenone | HMDB0033910 | 0.00016139 | 1.6115 | 0.002843 | 0.973517 |
| 7 | Methyl stearate | HMDB0034154 | 0.00066907 | 1.3375 | 0.006248 | 0.849138 |
| 8 | Urea | HMDB0000294 | 0.00088321 | 1.8096 | 0.007329 | 0.913793 |
| 9 | Phenethyl isocyanate | HMDB0256402 | 0.0011754 | 1.4531 | 0.008816 | 0.827586 |
| 10 | 2,5-pyrrolidinedione | HMDB0240653 | 0.0046311 | 1.5492 | 0.024742 | 0.857759 |
| 11 | Fumaric acid | HMDB0000134 | 0.0070368 | 2.6569 | 0.032192 | 0.857759 |
| 12 | 1,3-Propanediol | METPA0292 | 0.0075143 | 1.9584 | 0.032562 | 0.810345 |
| 13 | Scyllo-Inositol | HMDB0006088 | 0.0077449 | 1.0824 | 0.033192 | 0.902215 |

**Table S2:** List of significantly altered metabolites between COPD smokers and Healthy controls with VIP >1, p-value <0.05, FDR <0.05, AUC>0.8, and $Log_2FC \geq \pm 1$.

| S.No. | Metabolites | HMDB ID | p-value | VIP | $Log_2FC$ | FDR | AUC |
|---|---|---|---|---|---|---|---|
| 1 | Acetophenone | HMDB0033910 | 0.000008158 | 1.571996 | -3.0734 | 0.001615 | 1 |
| 2 | Pentadecane 2,6,10-trimethyl | | 0.000028553 | 1.127812 | -1.4447 | 0.001885 | 0.982 |
| 3 | Heptane, 3-methyl | HMDB0031583 | 0.000077501 | 1.626661 | -2.8554 | 0.003069 | 0.962333 |
| 4 | Fumaric acid | HMDB0000134 | 0.00018356 | 2.189606 | -3.3838 | 0.004543 | 0.951367 |
| 5 | Heptane 4-ethyl | | 0.000273 | 1.714466 | -2.4531 | 0.005411 | 0.933333 |
| 6 | Cyclopentanone | HMDB0031407 | 0.00027329 | 1.54909 | -2.3515 | 0.005411 | 0.952333 |
| 7 | Urea | HMDB0000294 | 0.00039159 | 1.78843 | -4.3001 | 0.007049 | 0.925 |
| 8 | Cyclopentane | | 0.00055475 | 1.474997 | -1.9887 | 0.009153 | 0.916667 |
| 9 | Arachidic acid | HMDB0002212 | 0.0010524 | 1.464422 | -2.3487 | 0.014884 | 0.9 |

| | | | | | | | |
|---|---|---|---|---|---|---|---|
| 10 | Phthalic acid, 6-ethyl, 3-octyl, butyl ester | | 0.0010524 | 1.449852 | -2.2494 | 0.014884 | 0.9 |
| 11 | 1,3-Propanediol | METPA0292 | 0.0024841 | 1.633006 | -1.8306 | 0.028933 | 0.875 |
| 12 | s-Triazine, 2-amino-4-(piperidinomethyl)-4-piperidino | | 0.0024841 | 1.515899 | -2.0465 | 0.028933 | 0.875 |
| 13 | 2-Pyrrolidinone | HMDB0002039 | 0.0032387 | 1.183363 | -1.9874 | 0.031747 | 0.866667 |
| 14 | Scyllo-Inositol | HMDB0006088 | 0.0032387 | 1.126073 | -1.7654 | 0.031747 | 0.866667 |
| 15 | Tetradecane, 4-methyl | HMDB0302457 | 0.0041688 | 1.112868 | -1.5451 | 0.031747 | 0.874333 |
| 16 | Dodecane | HMDB0031443 | 0.0041688 | 1.566589 | -1.5999 | 0.031747 | 0.858333 |
| 17 | Triethylene glycol | HMDB0259193 | 0.0041688 | 1.439016 | -1.5656 | 0.031747 | 0.858333 |
| 18 | Bis(2-isopropyl-5-methylcyclohexyl) methylphosphonate (isomer 1) | | 0.0041688 | 1.245875 | -1.3116 | 0.031747 | 0.858333 |
| 19 | Hexadecane | HMDB0033792 | 0.0041688 | 1.295447 | -1.5238 | 0.031747 | 0.858333 |
| 20 | Acetic acid | HMDB0000042 | 0.0041688 | 2.189344 | -2.3753 | 0.031747 | 0.858333 |
| 21 | Propiophenone 2-methyl | | 0.0067345 | 1.327379 | -2.2317 | 0.043014 | 0.851667 |
| 22 | 2,5-pyrrolidinedione | HMDB0240653 | 0.0067345 | 1.501022 | -3.4046 | 0.043014 | 0.841667 |
| 23 | Tetraethylene glycol | HMDB0094708 | 0.0067345 | 1.506442 | -2.1212 | 0.043014 | 0.841667 |
| 24 | Phosphoric acid dipentyl ethyl ester | | 0.0067345 | 1.215505 | -2.039 | 0.043014 | 0.841667 |
| 25 | Benzyl alcohol | HMDB0003119 | 0.0084558 | 1.414318 | -2.1729 | 0.049243 | 0.833333 |
| 26 | Undecane 5,7 dimethyl | | 0.0084558 | 1.865594 | -1.4355 | 0.049243 | 0.833333 |

**Table S3:** List of significantly altered metabolites between COPD Ex-smoker and Healthy controls having VIP >1, p-value <0.05, FDR <0.05, AUC>0.8, and $Log_2FC \geq \pm 1$.

| S. No. | Metabolites | HMDB ID | p-value | VIP | $Log_2FC$ | FDR | AUC |
|---|---|---|---|---|---|---|---|
| 1 | Glycolic acid | HMDB0000115 | 0.000026461 | 1.290376 | 1.7105 | 0.00116 | 1 |
| 2 | Cyclic octaatomic sulfur | HMDB0302183 | 0.000026461 | 2.257769 | -4.1194 | 0.00116 | 1 |
| 3 | Acetophenone | HMDB0033910 | 0.000026461 | 2.342066 | -4.884 | 0.00116 | 1 |
| 4 | Tetradecane, 4-methyl | HMDB0302457 | 0.000026461 | 2.937627 | -9.1263 | 0.00116 | 1 |
| 5 | Cyclopentanone | HMDB0031407 | 0.000026461 | 2.27208 | -5.455 | 0.00116 | 1 |

| | | | | | | | |
|---|---|---|---|---|---|---|---|
| 6 | Undecane, 2,4-dimethyl | | 0.000052923 | 2.26212 | -3.7108 | 0.00116 | 1 |
| 7 | Cyclopentane | | 0.000052923 | 2.180339 | -3.6887 | 0.00116 | 0.988636 |
| 8 | Propiophenone 2-methyl | | 0.000052923 | 2.066374 | -4.8393 | 0.00116 | 1 |
| 9 | 4,4 dimethyl octane | | 0.000052923 | 2.439377 | -8.5553 | 0.00116 | 0.988636 |
| 10 | N-Acetyl glucosamine | HMDB0000803 | 0.000106 | 1.84769 | 3.6833 | 0.001885 | 0.981173 |
| 11 | Diethylene glycol adipate | | 0.00010585 | 2.340317 | -7.2585 | 0.001885 | 0.977273 |
| 12 | 2-sec-Butyl-6-ethylaniline | | 0.00018523 | 1.742163 | -2.5134 | 0.002933 | 0.965909 |
| 13 | Hydracrylic acid | HMDB0000700 | 0.000318 | 1.635309 | 3.1849 | 0.004525 | 0.954545 |
| 14 | 3-Hydroxybutyric acid | HMDB0000011 | 0.000794 | 1.297349 | 1.9865 | 0.008702 | 0.931818 |
| 15 | D-(+)-Talose | | 0.000794 | 1.48408 | 2.5727 | 0.008702 | 0.931818 |
| 16 | Ethylparaben, acetate | HMDB0032573 | 0.0011908 | 1.563142 | -1.7746 | 0.010284 | 0.920455 |
| 17 | Scyllo-Inositol | HMDB0006088 | 0.0011908 | 1.68342 | -2.3626 | 0.010284 | 0.930455 |
| 18 | 4-tert-Octylphenol | HMDB0013825 | 0.0011908 | 1.946461 | -3.6939 | 0.010284 | 0.920455 |
| 19 | Hexadecanoic acid, ethyl ester | HMDB0029811 | 0.0011908 | 1.936855 | -2.274 | 0.010284 | 0.920455 |
| 20 | Phenethyl isocyanate | HMDB0256402 | 0.0011908 | 1.823966 | -3.3314 | 0.010284 | 0.920455 |
| 21 | Methyl stearate | HMDB0034154 | 0.0011908 | 2.006405 | -2.6024 | 0.010284 | 0.920455 |
| 22 | Acenaphthene | HMDB0247899 | 0.0025403 | 1.080107 | -1.4666 | 0.020111 | 0.897727 |
| 23 | Bis(2-hydroxyethyl) phthalate | | 0.0035987 | 1.772683 | -1.803 | 0.025016 | 0.886364 |
| 24 | Thiophene | HMDB0029718 | 0.0035987 | 1.425036 | -2.9172 | 0.025016 | 0.886364 |
| 25 | Undecane, 2,6-dimethyl | | 0.003599 | 1.425647 | -1.6612 | 0.025016 | 0.886364 |
| 26 | Benzene 1,2,3 trimethyl | HMDB0059901 | 0.0035987 | 1.839217 | -1.6979 | 0.025016 | 0.886364 |
| 27 | 4-Hydroxybenzeneacetic acid | HMDB0000020 | 0.004975 | 1.444908 | 4.6873 | 0.031507 | 0.875 |

| | | | | | | | |
|---|---|---|---|---|---|---|---|
| 28 | 3,4-Dihydroxymandelic acid | HMDB0001866 | 0.0049747 | 1.74765 | -2.9372 | 0.031507 | 0.875 |
| 29 | 2,3-Butanediol | HMDB0003156 | 0.006801 | 1.34172 | 2.3297 | 0.038763 | 0.863636 |
| 30 | Hexadecane | HMDB0033792 | 0.009103 | 1.364083 | -1.1659 | 0.047169 | 0.852273 |
| 31 | Benzyl alcohol | HMDB0003119 | 0.009103 | 1.328314 | -1.6098 | 0.047169 | 0.852273 |

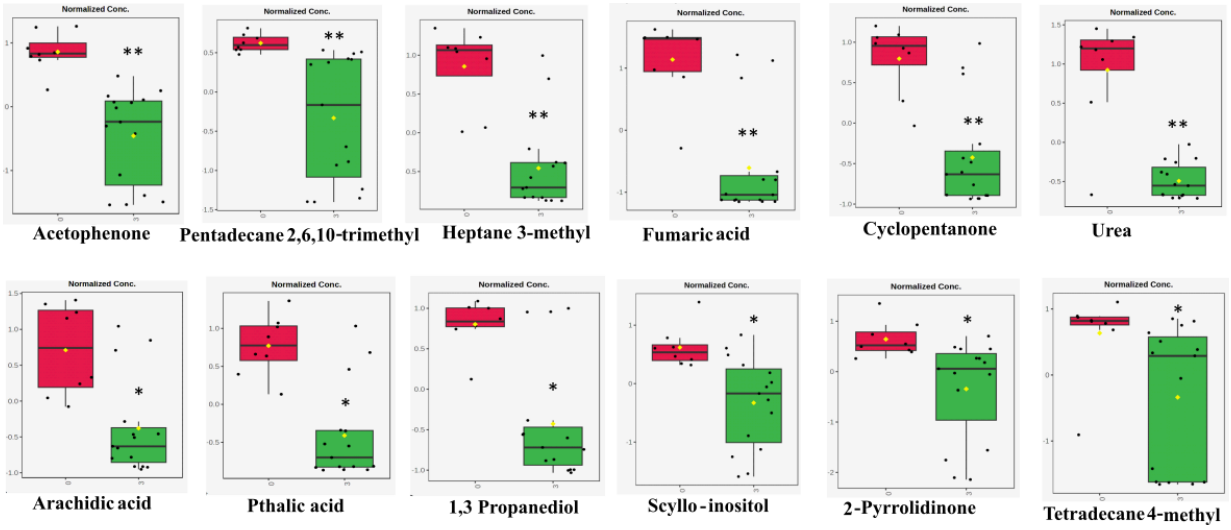


**Fig S1.** Box and whisker plots of the top 12 differentially expressed metabolites categorized based on the p-value in COPD smokers (Green) vs healthy controls (Red), with * denotes p< 0.05, **denotes p< 0.01.

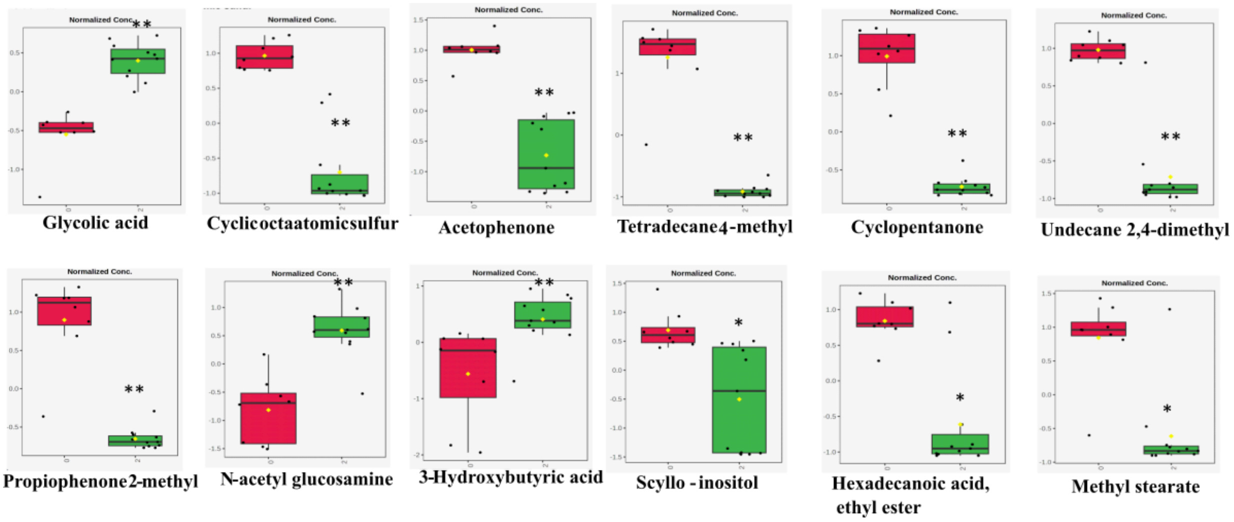


**Fig S2.** Box and whisker plots of the top 12 differentially expressed metabolites categorized based on the p-value in COPD ex-smoker (Green) vs healthy control (Red), with * denotes p-value < 0.05, and ** represents p-value < 0.01.

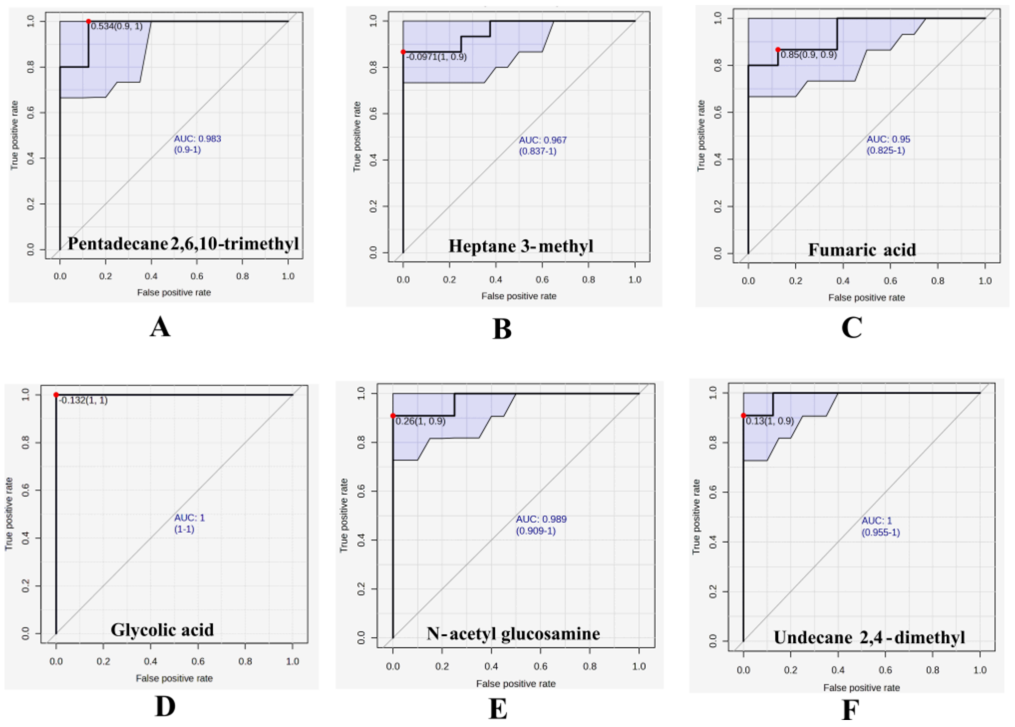


**Fig S3.** The receiver operating characteristic (ROC) curve for metabolites with an area under the curve (AUC) of ≥ 0.85 and a p-value of < 0.05 is presented for comparisons between COPD smokers and healthy controls (A-C), as well as COPD ex-smokers and healthy controls (D-F).

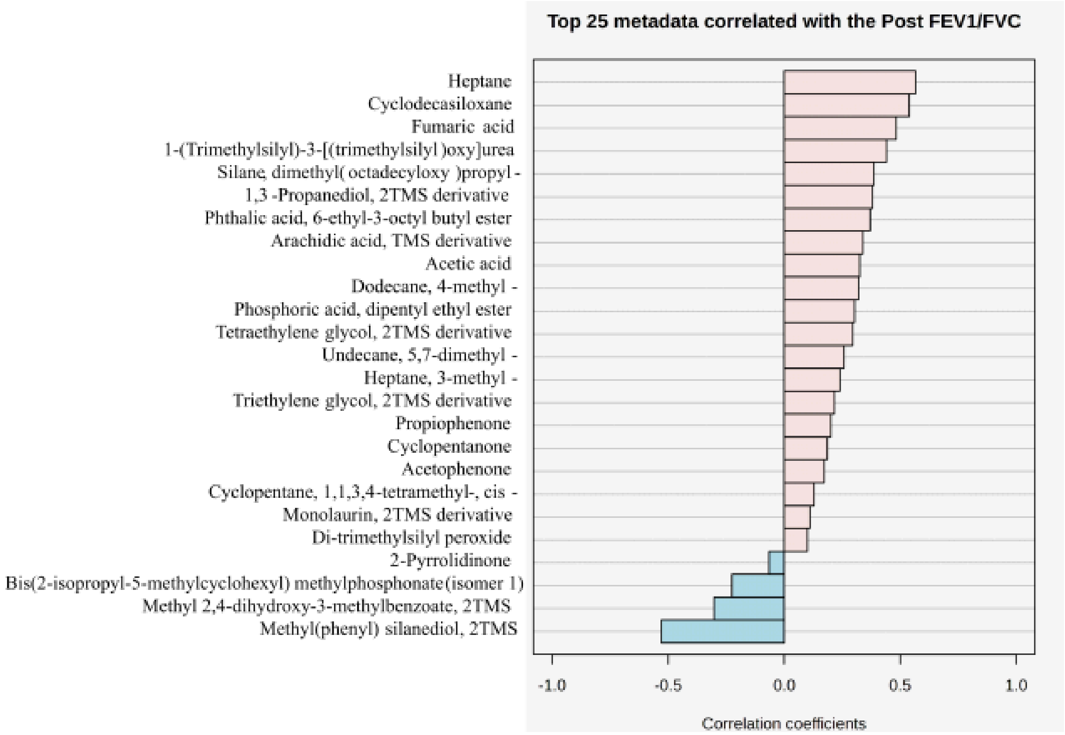


**Fig S4.** Spearman correlation plot of metabolites with post $FEV_1/FVC$ ratio of COPD smokers subgroup, with $\rho \geq 0.3$ and $p < 0.05$ considered a strong correlation. ($\rho$ is the correlation coefficient).

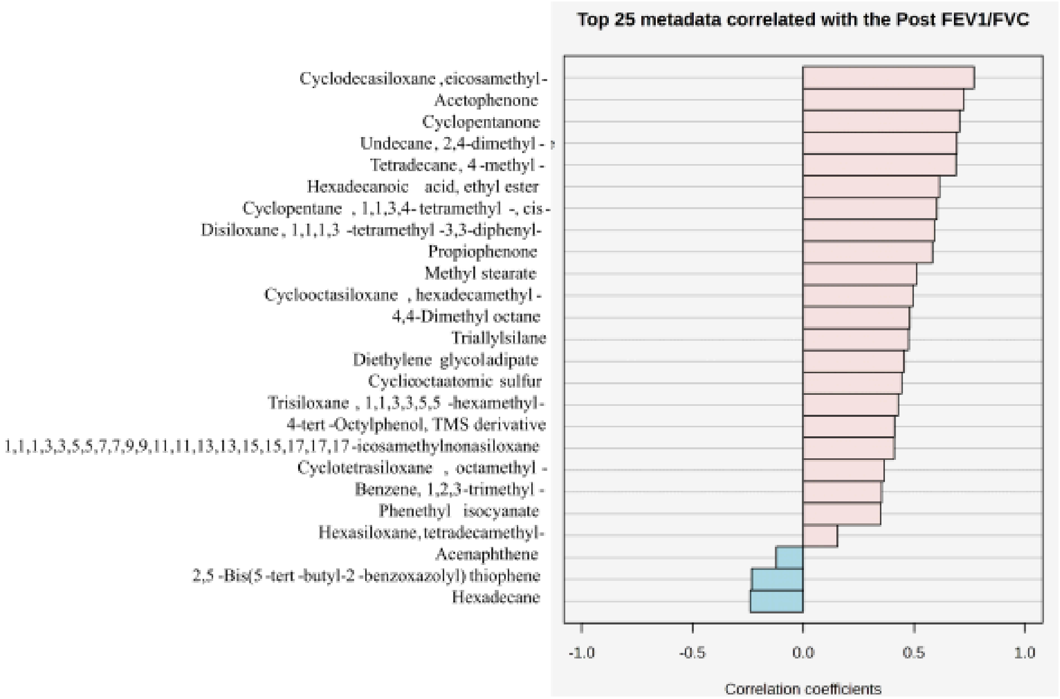


**Fig S5.** Spearman correlation plot of metabolites with post $FEV_1/FVC$ ratio of COPD Ex-smoker subgroup, with $\rho \geq 0.3$ and $p < 0.05$ considered a strong correlation. ($\rho$ is the correlation coefficient)

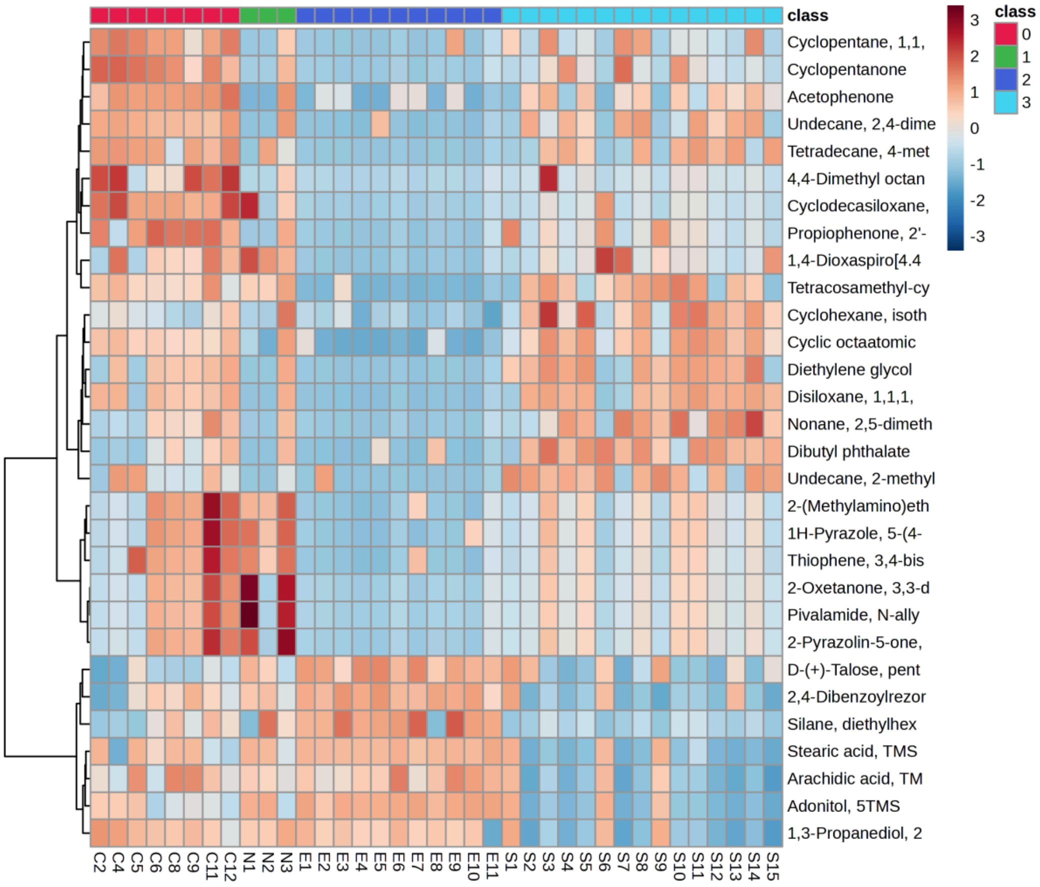


**Fig S6.** Heatmap depicting hierarchical clustering of differentially expressed metabolites across COPD subgroups and healthy controls with p-value < 0.05 (class 0 - Healthy control, class 1- Non-smoker, class 2- Ex-smoker, class 3- Smoker). The red-colored bars show upregulated metabolites, and the blue-colored bars represent downregulated metabolites.

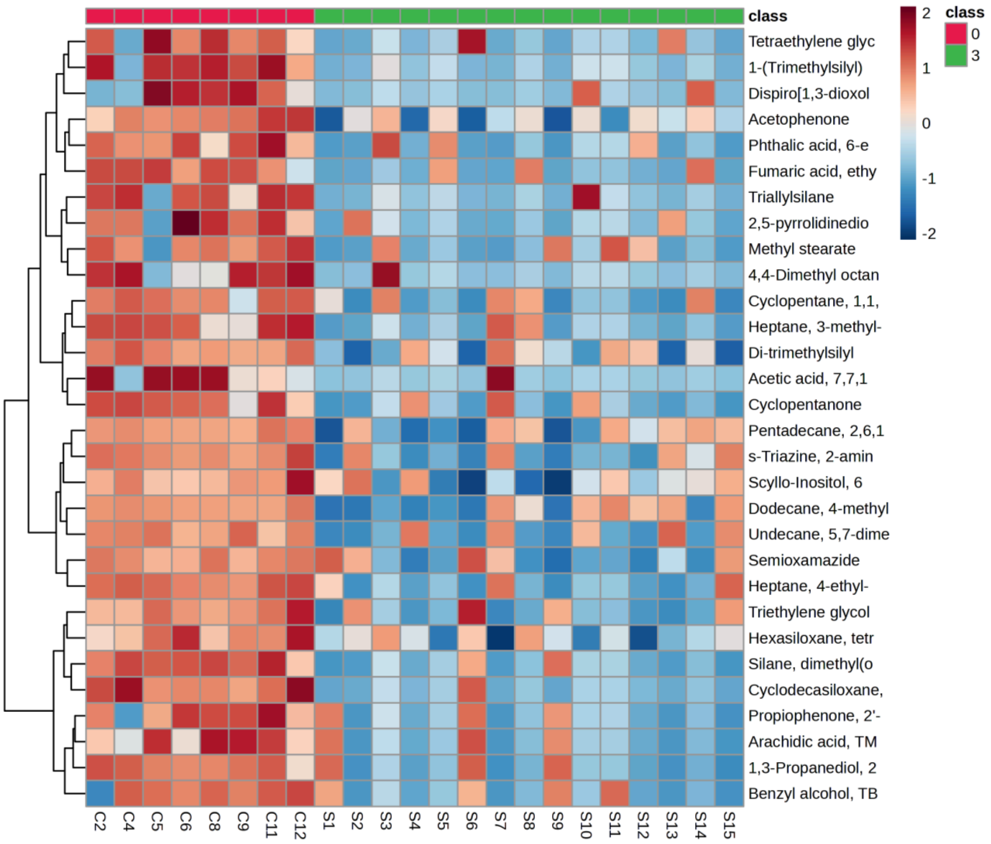


**Fig S7.** Heatmap depicting hierarchical clustering with differentially expressed metabolites of COPD smoker subgroups and healthy controls with p-value < 0.05 (class 0- Healthy control, class 3- Smoker). The red-colored bars show upregulated metabolites, and the blue-colored bars represent downregulated metabolites.

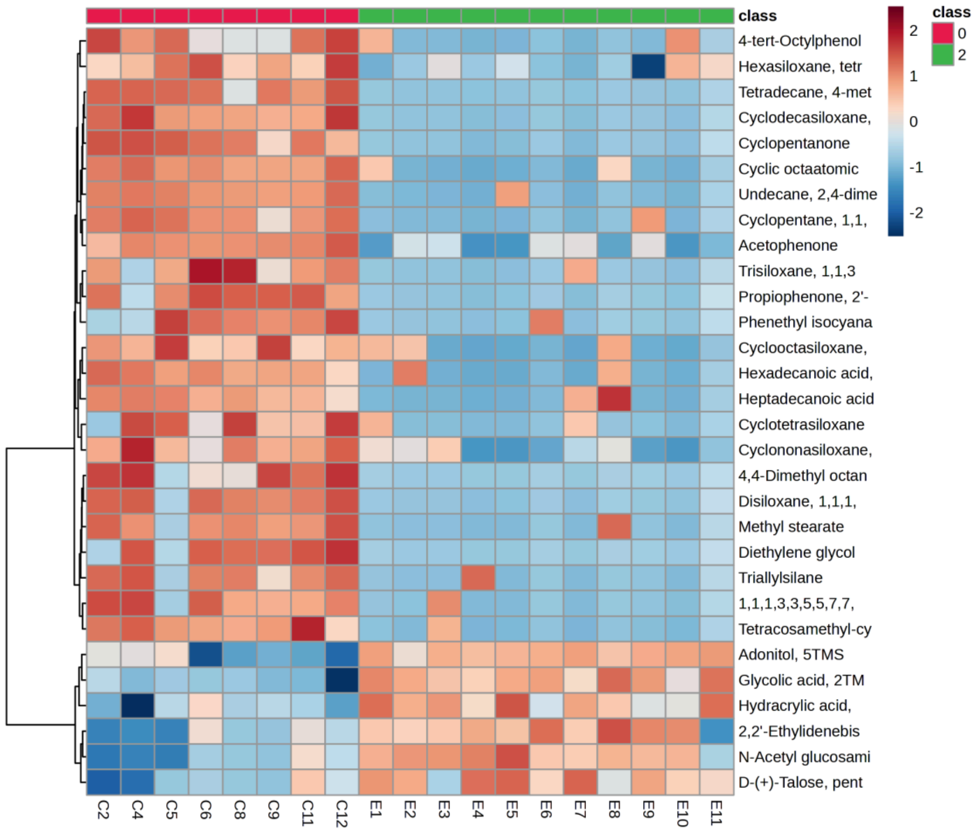


**Fig S8.** Heatmap depicting hierarchical clustering of differentially expressed metabolites in COPD ex-smoker subgroups and healthy controls with p-value < 0.05 (class 0- Healthy control, class 2- Ex-smoker). The red-colored bars show upregulated metabolites, and the blue-colored bars represent downregulated metabolites.
